# Two chromosome-level genome assemblies of *Sarracenia* reveal repeat-driven expansion and gene loss associated with carnivory

**DOI:** 10.64898/2026.03.01.708852

**Authors:** Ethan A. Baldwin, Willie L. Rogers, Jim Leebens-Mack

**Affiliations:** Department of Plant Biology, University of Georgia, Athens, GA, United States

**Keywords:** carnivory, genome assembly, gene family evolution, gene loss, photosynthesis, comparative genomics, *Sarracenia*

## Abstract

**Premise of the Study:** Carnivory has evolved repeatedly across the plant tree of life despite being a dramatic shift from typical plant nutrient acquisition strategies. It remains largely unclear whether the evolution of carnivory takes a similar genomic trajectory. Here, we explore the genomic consequences of carnivory in the pitcher plant genus *Sarracenia*.

**Methods:** We use a combination of Pacbio HiFi long-read sequencing and trio-binning to assemble chromosome-scale genome sequences for *S. psittacina* and *S. rosea*. We conduct comparative analyses with other asterid genomes to evaluate patterns of gene family expansion and contraction during the transition to carnivory.

**Results:** Both *Sarracenia* genomes are large (∼3.5 Gbp) and highly repetitive (∼87% repeats) yet only contain ∼22,000 genes. This reduced gene content reflects widespread gene family contraction. In total, 3,654 gene families have contracted, including the complete loss of 934 gene families, while only 751 gene families have expanded. The gene losses are enriched for functions related to photosynthesis, including nuclear-encoded subunits of the NADH dehydrogenase (Ndh) complex, as well as immune-related genes.

**Conclusions:** These results indicate that the evolution of carnivory in *Sarracenia* is associated with widespread gene loss rather than extensive gene family expansion. The loss of genes involved in photosynthesis and immune response suggest the relaxation of selection on these functions, which may be partially supplanted by prey-derived nutrient acquisition and pitcher-associated microbiome. These chromosome-level assemblies will enable future comparative studies in plant evolution, while also serving as critical resources for the conservation of this ecologically significant lineage.

## INTRODUCTION

Carnivorous plants capture and digest insect prey to acquire nutrients. The evolution of carnivory requires the emergence of specialized structures that attract and trap prey, along with the physiological mechanisms to digest prey and absorb released nutrients. Remarkably, this complex suite of traits has arisen at least 10 times across the angiosperm tree of life (Fleischmann et al. 2017). The repeated evolution of carnivory has long been studied as a model for the convergent evolution of complex traits (Albert et al. 1992), yet the genetic basis of these adaptations remain largely unclear. In particular, it is not known whether the repeated emergence of carnivory is accompanied by shared genomic changes or if lineages take unique evolutionary routes towards carnivory.

High-quality genome assemblies can enable research aimed at answering these questions, however reference-quality genome assemblies and annotations have been published for only four of the ten carnivorous lineages (Fukushima et al. 2017;Lan et al. 2017;Hartmann et al. 2020;Palfalvi et al. 2020). Of these, only the Lentibulariaceae (bladderwort) and Caryophylales (venus fly trap and pitcher plants in the genus *Nepenthes*) reference genomes have been assembled using long-read sequencing technologies, which produce genomes that are significantly more complete and contiguous than short-read assemblies (Paajanen et al. 2019). Expanding genomic resources to other lineages, including the pitcher plant family Sarraceniaceae, is essential for understanding the genomic basis of carnivory and its repeated evolution.

The pitcher plant genus *Sarracenia* L. (Sarraceniaceae, Ericales) comprises charismatic carnivorous plants native to eastern North America. Each of the 10 species of *Sarracenia* produces tube-shaped leaves that attract, capture, and digest insect prey, providing the plant with mineral nutrients. *Sarracenia* species perform vital ecosystem services, providing moist microenvironments for numerous inquiline arthropod species (Harvey and Miller 1996) and serving as host for moths in the genus *Exyra* (Noctuidae), which spend most of their lives in *Sarracenia* pitchers (Stephens and Folkerts 2012). Due to widespread loss of *Sarracenia*’s habitat, almost all species are of conservation concern, including two taxa that are federally endangered (*S. jonesii* and *S. oreophila*). With no genome assemblies available for *Sarracenia*, population genetic studies of these species have been restricted to a few microsatellite loci, severely limiting the ability of evolutionary biologists and conservation geneticists to survey genetic diversity within and between populations (Koopman and Carstens 2010;Furches et al. 2013;Rentsch and Holland 2020).

Here, we exploit recent technological advances to generate fully phased chromosome-level genome assemblies from two *Sarracenia* species from PacBio HiFi and Omni-C reads derived from an F1 hybrid. We use the reference genome assemblies to infer ancestral shifts in gene content that we hypothesize are associated with the evolution of carnivory. In addition, these assemblies will serve as invaluable resources for conservation practitioners working to safeguard the remaining populations of rare *Sarracenia* species. They will also be integral to broader comparative analyses seeking to understand the mechanisms underlying the repeated evolution of carnivory in angiosperms.

## MATERIALS AND METHODS

### DNA and RNA extraction and sequencing

*Sarracenia rosea* has pitchers with wide openings that are typical for *Sarracenia* species and other pitcher plants with pitfall traps, while *S. psittacina* has pitchers with extremely narrow, funnel-shaped openings which act as “lobster pot traps” (Fig. 1A). *S. psittacina, S. rosea*, and their F1 hybrid were obtained from a greenhouse at UGA, where clones have been maintained for nearly two decades (Malmberg et al. 2018). High molecular weight DNA was obtained from the F1 by first isolating nuclei according to the “Isolating nuclei from plant tissue using TissueRuptor disruption” protocol available from PacBio, and then extracting the DNA from the nuclei using the Nanobind plant nuclei kit. The high molecular weight DNA was sent to Hudson Alpha where SMRTbell libraries were prepared and sequenced on four SMRT cells on a Revio. Tissue from the F1 was sent to Hudson Alpha where an Omni-C library was prepared and sequenced on an S4 flow cell on a NovaSeq 6000.

**Figure 1.**
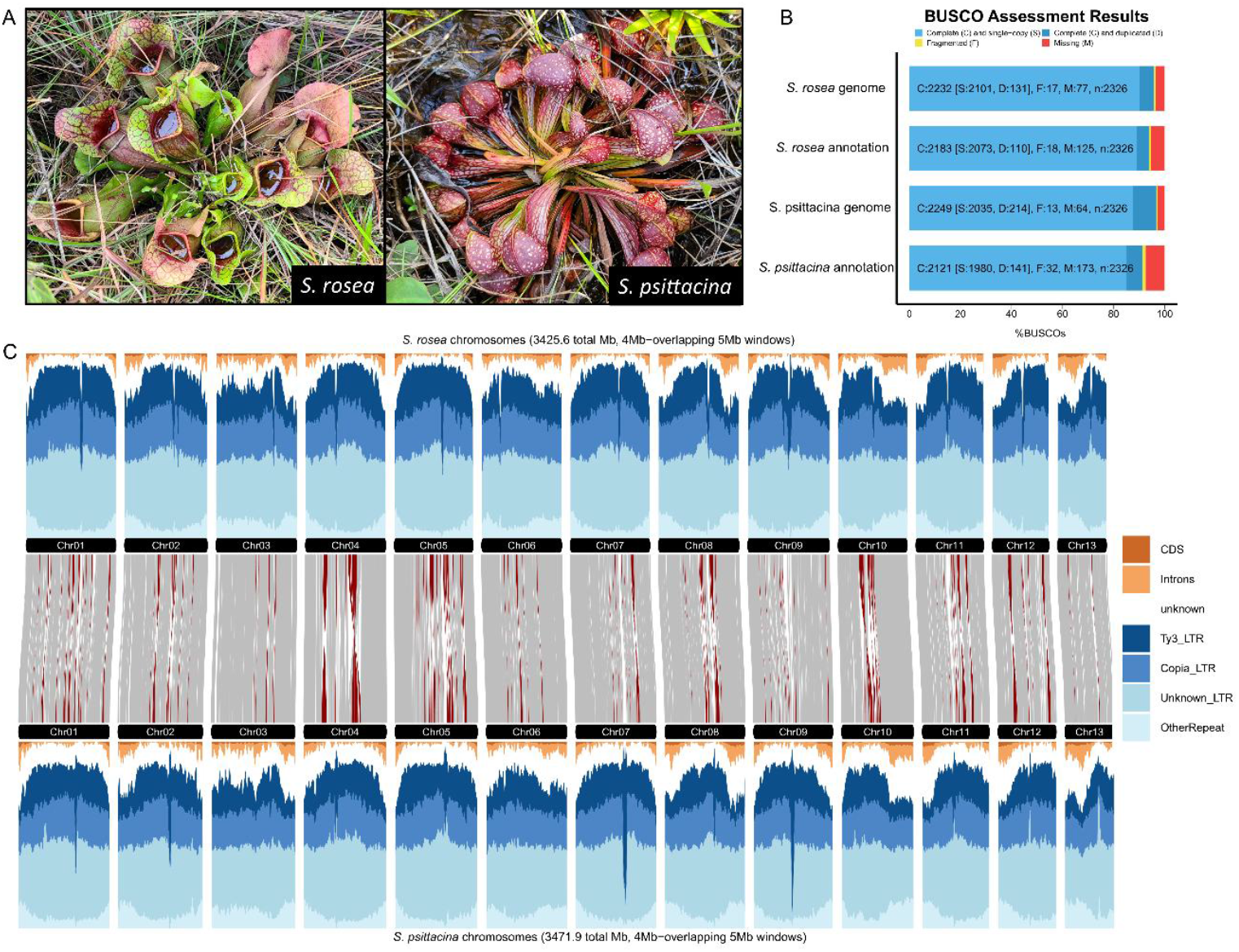
*Sarracenia* and its genome assembly. (A) In-situ photographs of *S. rosea* and *S. psittacina*. (B) BUSCO results. (C) Comparison of gene and repeat content between *S. rosea* and *S. psittacina* genome assemblies. Grey bars between the two genomes represent collinear syntenic blocks, while rearrangements are represented in red.

Illumina shotgun sequencing was performed on the parents of the sequenced F1 genotype for trio-binning (Koren et al. 2018). DNA was isolated from fresh or dry tissue using Qiagen DNeasy Plant Pro kits. Libraries were constructed with Kapa HyperPlus library kits using custom adapters and iTru primers from Adapterama I (Glenn et al. 2019). Sequencing was done on 10B flow cells on a NovaSeq X at SeqCenter in Pittsburgh, PA.

RNA was extracted from *S. psittacina* and *S. rosea* young pitchers, mature pitchers, and roots using Zymo Direct-zol RNA kits, substituting Invitrogren Plant RNA reagent for the TRI reagent. Sequencing libraries were constructed using Kapa Stranded mRNA-Seq kits using custom adapters and iTru primers from Adapterama I (Glenn et al. 2019). Sequencing was done on 10B flow cells on a NovaSeq X at SeqCenter in Pittsburgh, PA.

### Genome assembly and scaffolding

Assembly of the F1 HiFi reads was performed with the trio binning (Koren et al. 2018) method in hifiasm v. 0.19.8 (Cheng et al. 2021). Parental 31-mers were counted from the shotgun sequencing reads using yak (https://github.com/lh3/yak) and used as input for hifiasm.

Scaffolding was done with Omni-C sequencing data derived from the F1. The Omni-C reads were mapped to each assembly using BWA-MEM v. 0.7.17 (Li and Durbin 2009) with the-5SP flag. PCR duplicates were removed using SAMBLASTER v. 0.1.26 (Faust and Hall 2014) and reads were sorted using SAMtools v. 1.16.1 (Li et al. 2009). Initial scaffolding was done with YaHS v. 1.1 (Zhou et al. 2023), and final scaffolding was manually performed using Juicebox Assembly Tools v. 1.8.8 (Durand et al. 2016;Dudchenko et al. 2018). The completeness of the assemblies were assessed by scanning for conserved Eudicot single-copy orthologs using BUSCO v. 5.5.0 (Simão et al. 2015) with the Eudicot OrthoDB v. 11 database (Kuznetsov et al. 2022).

### Annotation

Repeat sequences in the genome assemblies were identified by using RepeatModeler v. 2.0.4 to produce TE libraries (http://www.repeatmasker.org/). The TE libraries were then used to annotate TEs and mask repeat regions using RepeatMasker v. 4.1.5 (http://www.repeatmasker.org/). The BRAKER3 pipeline v. 3.0.8 (Gabriel et al. 2024) was used to annotate protein coding genes using RNA-seq and protein homology. RNA-seq reads from *S. psittacina* and *S. rosea* were mapped to their respective genomes using HISAT v. 2.2.1 (Kim et al. 2019). Eudicot protein sequences were downloaded from OrthoDB v. 11 (Kuznetsov et al. 2022).

### Gene loss and gene family evolution analysis

Gene models for both parental haplotype assembles were assigned to orthogroups using OrthoFinder 2.5.4 (Emms and Kelly 2019) in an analysis including proteomes from nine other angiosperm species selected to represent major flowering plant clades and include multiple close relatives of *Sarracenia*. The additional species include *Amborella trichopoda*, the sister lineage to all other extant angiosperms, *Oryza sativa*, a monocot, *Arabidopsis thaliana*, the model asterids *Mimulus guttatus* and *Solanum lycopersicum*, and four species more closely related to *Sarracenia* within *Ericales – Vaccinium darrowii, Camellia sinensis, Actinidia eriantha*, and *Actinidia chinensis*. After orthogroup assignment, we analyzed the functional profile of orthogroups that were lost in *Sarracenia*. Lost orthogroups were defined as those where both *Sarracenia* species had no orthologs and at least four of the six rosid species contained orthologs. GO term enrichment of missing orthogroups was done using the clusterProfiler R package (Yu et al. 2012). An *Arabidopsis* ortholog from each of the missing orthogroups was used as the foreground set, and an *Arabidopsis* ortholog from each orthogroup was used as the background. P values were adjusted using false discovery rate, and a cut-off of 0.05 was used.

To model gene family evolution, we first generated a species tree with orthogroups identified as being conserved in single copy by OrthoFinder. To ensure accurate gene trees, OrthoFinder was run with the multiple sequence alignment option, and IQ-TREE2 was used as the gene tree estimation software (Nguyen et al. 2015). The species tree output from OrthoFinder was made ultrametric using the make_ultrametric.py command in OrthoFinder with a root age of 140 million years ago based on the divergence time between *Amborella* and all other angiosperms (Magallón et al. 2015). To reduce the complexity of the model, the species tree was pruned to include only the Ericales genomes and *Solanum* as an outgroup. CAFE5 (Mendes et al. 2020) was used to model gene family expansion and contraction along this reduced species tree under the gamma model with the number of rate categories (k) varying from 1-4 to determine which k value best fit the data. A k value of 3 was found to have the highest likelihood, so all results presented are from the k=3 model. Orthogroups with more than 100 orthologs in any given species were removed from the analysis using the clade_and_size_filter.py script from CAFE. The functional enrichment of expanding and contracting gene families on the branch leading to *Sarracenia* were analyzed for GO term enrichment as described above.

## RESULTS

### Two chromosome scale reference genomes for *Sarracenia*

We assembled *S. rosea* and *S. psittacina* genomes using 337gb of Pacbio Hifi reads from their F1 (∼48X coverage per haplotype) in addition to Illumina WGS reads from both parents for trio binning. The resulting genome assemblies are 3488mb and 3594mb long for S. rosea and S. psittacina respectively, coinciding closely with published estimates of genome sizes across Sarracenia based on flow cytometry (Veleba et al. 2020). After scaffolding using Omni-C reads generated from the F1, 96.6% of *S. psittacina* and 98.2% of *S. rosea* assemblies were assigned to psuedochromosomes (Table 1). Approximately 22000 gene models were annotated in each of the assemblies and the embryophyte BUSCO scores of 97.8% and 98.9% for *S. psittacina* and *S. rosea* respectively (Fig. 1B) (Table 1).

**Table 1.** Genome assembly statistics.

|  | *S. rosea* | *S. psittacina* |
| --- | --- | --- |
| Total assembly length | 3488 mb | 3594 mb |
| Chromosome length | 3426 mb | 3472 mb |
| Small scaffold length | 62 mb | 122 mb |
| Scaffold N50 | 280 mb | 281 mb |
| Contigs N50 | 9 mb | 7 mb |
| Number of scaffolds | 896 | 1161 |
| Number of contigs | 1505 | 1847 |
| Number of genes | 21900 | 22411 |
| Complete BUSCOs | 1564 | 1579 |
| Complete and single-copy BUSCOs | 1494 | 1451 |
| Complete and duplicated BUSCOs | 70 | 128 |
| Fragmented BUSCOs | 14 | 8 |
| Missing BUSCOs | 36 | 27 |
| Total BUSCO groups searched | 1614 | 1614 |

Both *Sarracenia* genomes were found to be highly repetitive, with annotated repeats comprising ∼87% of the assemblies, which is considerably higher than any other sequenced genome in the Ericales (*Diospyros oleifera* is next highest at 64.96%) (Zhu et al. 2019). The most abundant repetitive elements are LTR retrotransposons, half of which belong to unknown LTR families and the remainder approximately equal proportions of Ty1 and Ty3 elements. These elements are concentrated in pericentromeric regions, while genes are concentrated in the chromosome arms (Fig. 1C), which is a typical configuration for angiosperm genomes (Neumann et al. 2011;Sigman and Slotkin 2016). *S. rosea* and *S. psittacina* genomes are highly colinear across gene-rich regions, with many small structural rearrangements in the pericentromeric regions (Fig. 1C).

### Significant gene loss in *Sarracenia*

Despite its large genome size and high percentage of retained BUSCO genes (Fig 1B), *Sarracenia* has significantly fewer annotated gene models than its relatives in the Ericales due to widespread gene family contractions and the complete loss of many gene families following divergence from other lineages in the order (Fig 2). We found that 3654 gene families have contracted on the branch leading to the last common ancestor of the two *Sarracenia* species while only 751 gene families have expanded. This is by far the largest number of contracted gene families in any of the internal branches in this analysis. Interestingly, the terminal branch leading to *Camellia* has more gene family contractions than seen on the branch leading to *Sarracenia*, but it also has a large number of gene family expansions (Fig 2). In addition, 934 gene families that are conserved in the six other asterids included in our analysis are completely absent in the *Sarracenia* genomes.

**Figure 2.**
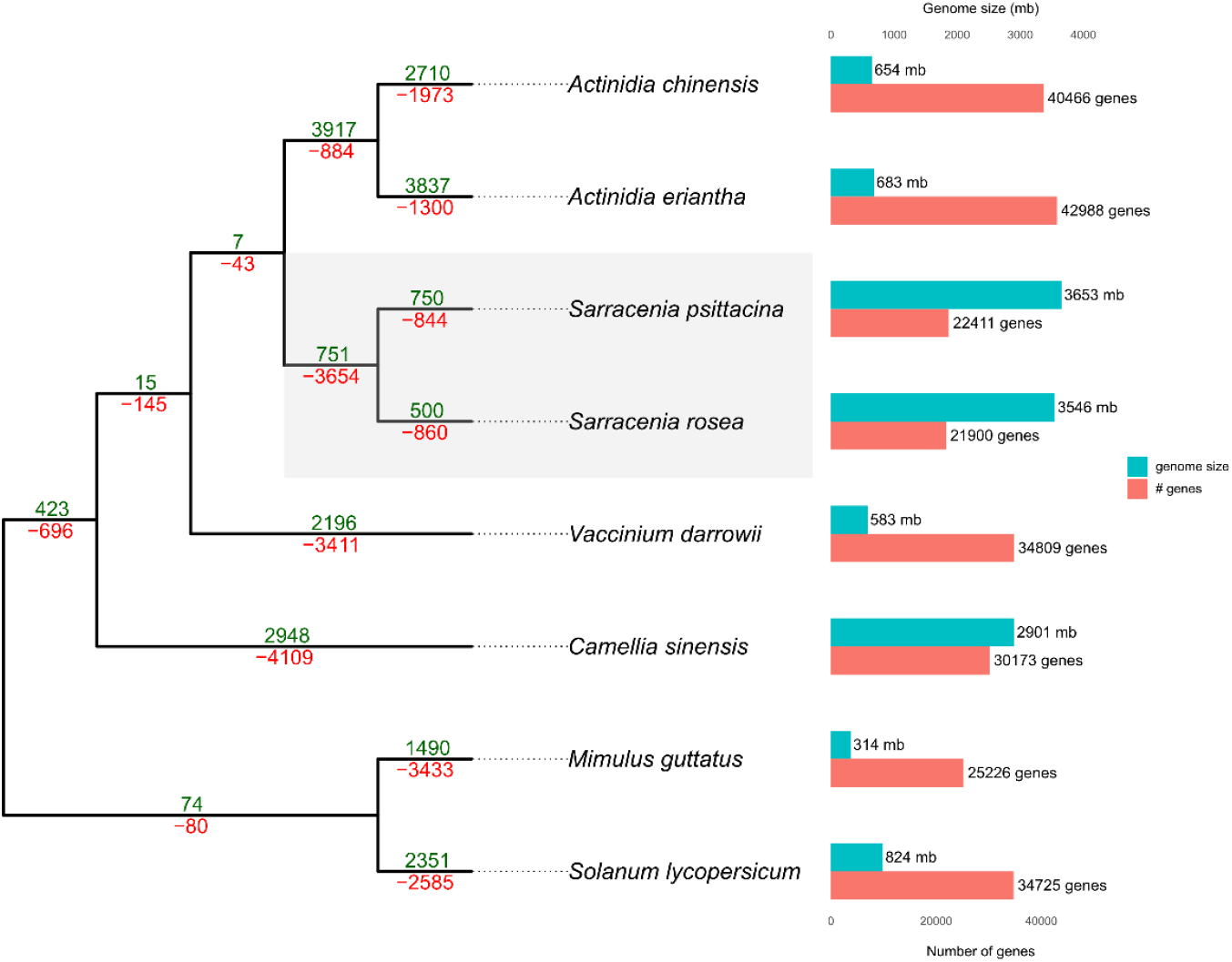
Summary of orthogroup expansion and contraction. Branches are labeled with the number of orthogroups that are either expanding (green, above) or contracting (red, below) on that branch. Aligned with the tips of the tree are bar plots of genome size and number of annotated gene models in each genome.

### Loss of photosynthesis and immune response genes

GO term enrichment of the contracted and missing gene families in *Sarracenia* genomes revealed the loss of genes related to several key biological functions. Whereas just one GO functional annotation term (translation) was enriched for gene families that expanded on the branch leading to *Sarracenia*, 3654 contracted and 429 lost gene families were enriched in several key biological functions relating to photosynthesis (Fig. 3). The enrichment for loss of photosynthesis-related genes is primarily driven by the absence of the majority of genes involved in the NADH dehydrogenase (*Ndh*) complex assembly (Fig. 3 B,D). While *Ndh* genes are not strictly necessary for photosynthetic function, the *Ndh* complex plays a role in photosynthetic electron transport and is important for maintaining photosynthetic efficiency under certain environmental stresses (Graham et al. 2017). Many of the plastome-encoded *Ndh* subunits are absent or psuedogenized in *Sarracenia* plastomes (Baldwin et al. 2023) and other carnivorous plant plastomes (Fu et al. 2023), but this is the first case where the loss of nuclear-encoded *Ndh* complex and related genes has been shown. The loss of photosynthesis genes is specific to *Sarracenia* and not seen in the other Ericales genomes (Fig. 3E).

**Figure 3.**
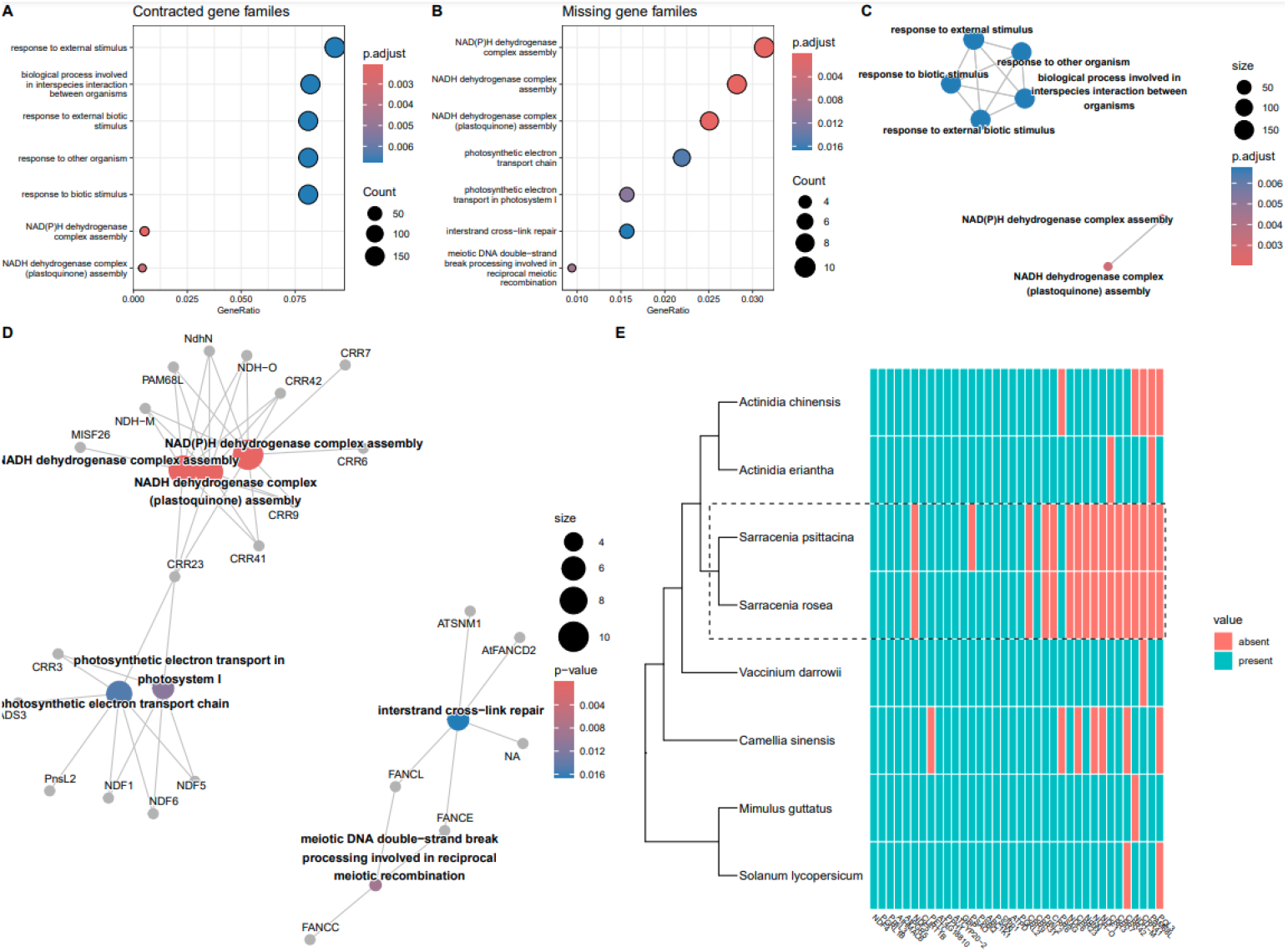
Functional enrichment of missing genes in *Sarracenia* genomes. (A) GO term enrichment of contracted gene families and (B) missing gene families. (C) Enrichment map of contracted gene families. GO terms are connected by edges when gene sets are overlapping. (D) Gene concept network for missing gene families. Enriched terms are hubs and the genes annotated with those terms are connected to them by edges. (E) Presence/absence of all genes in the 5 enriched photosynthesis-related GO terms.

The contracted orthogroups are also enriched in a cluster of GO terms related to responses to other organisms (Fig. 3A,C). Many of the contracted orthogroups associated with those GO terms are involved in immune response. For example, *Sarracenia* does not have any genes in the orthogroup containing members of the receptor-like protein (RLP) superfamily, while its close relatives in the Ericales have >10 RLP copies. RLPs are transmembrane receptors typically involved in the detection of pathogenic microbes and signaling an immune response (Jamieson et al. 2018). Other orthogroups where *Sarracenia* has significant contractions include most of the NB-ARC domain containing orthogroups, which are also known to play a role in pathogen resistance (van Ooijen et al. 2008).

## DISCUSSION

By leveraging a combination of state-of-the-art sequencing technologies and the trio-binning technique, we have produced chromosome-level genome assemblies for two distinct *Sarracenia* species. These are the first published genome assemblies for carnivorous plants in the Sarraceniaceae and the first long-read assemblies published for any carnivorous member of the Ericales, although a short read assembly has been published for *Roridula* (Hartmann et al. 2020). These high-quality assemblies have broadened the representation of carnivorous plant lineages with available genomic resources, and will be critical resources for identifying genomic changes associated with the evolution of carnivory.

*Sarracenia* species have large genomes filled with repetitive content, but there has been significant gene loss possibly associated with the evolution of carnivory. This architecture closely parallels that of the venus flytrap (*Dionaea muscipula*) genome. While close carnivorous relatives of *D. muscipula* exhibit less extreme genome sizes and repeat content, their genomes exhibit similar reductions in gene number (Palfalvi et al. 2020). *Utricularia gibba*, another carnivorous plant, exhibits an inverse pattern. *Utricularia* has a gene count that is more typical of flowering plants, but its genome size is the smallest of all vascular plants (Ibarra-Laclette et al. 2013). These results suggest that carnivory does not have a predictable effect on plant genome size, repeat content, or gene number, despite carnivorous plants containing several outliers.

The reduced number of genes in the *Sarracenia* genomes analyzed here is the result of widespread gene family contraction, including the complete loss of many gene families. Significant gene loss has previously been associated with a shift to parasitism (Sun et al. 2018;Cai et al. 2021), primarily driven by a relaxation of selection for traits no longer necessary for a parasitic lifestyle, such as photosynthesis. While carnivorous plants have been reported to have lost photosynthetic genes in their plastid genomes —namely those that code for subunits of the *Ndh* complex (Wicke et al. 2013;Nevill et al. 2019;Baldwin et al. 2023)—we are the first to report the loss of photosynthesis-related genes in the nuclear genome of a carnivorous plant. Carnivorous plants are well-known for their ability to obtain nitrogen and phosphorous from their digested prey, but the dispensability of photosynthesis-related genes could be indicative of assimilation of carbon from prey and a resulting relaxation of the need for efficient photosynthetic machinery. A few studies have indicated that carnivorous plants are obtaining some carbon from their prey (Kruse et al. 2014;Fasbender et al. 2017;Kruse et al. 2017;Pavlovič 2022;Lin et al. 2025), but the loss of *Ndh* genes in both the plastid and nuclear genomes of *Sarracenia* warrants further study on the functional impact of prey-derived carbon.

While the loss of nuclear *Ndh* genes may hint at a relaxation of selection for efficient photosynthesis, we hypothesize that the loss of immune response genes is an adaptation associated with *Sarracenia*’s mode of prey capture and nutrient absorption. *Sarracenia*’s pitchers contain decomposing insect prey in addition to a diverse assemblage of bacteria, fungi, and other organisms (Ellison et al. 2003). The microbial component of this community are important for prey digestion, potentially even more important than enzymes secreted by the pitcher itself (Luciano and Newell 2017). Distinctive bacterial communities have been described in different sub-habitats of pitchers, including embedded within the wall of the pitcher themselves. An immune response in *Sarracenia* pitchers may negatively impact the beneficial microbial community or reduce the transfer of nutrients from decomposing microbes into absorptive *Sarracenia* cells. Unlike most other carnivorous plant genomes, we do not see an expansion of digestive enzyme gene families in *Sarracenia*. We hypothesize that *Sarracenia* may rely on their symbiotic microbial community for the digestion of insect prey than other carnivorous plants with gene repertoires enabling production of a broader array of digestive enzymes.

## CONCLUSIONS

We present here two high-quality chromosome-scale genome assemblies from the pitcher plant species *Sarracenia psittacina* and *Sarracenia rosea*. These are the first published reference genomes for the carnivorous plant family Sarraceniaceae, filling a critical gap in the genomic resources available for carnivorous plants. We show that carnivory has left a unique imprint on the genome of *Sarracenia*, reflecting an evolutionary trajectory that is distinct from that of other carnivorous lineages. Additionally, the genomic resources presented here will enable more powerful analyses of genetic variation within and among *Sarracenia* taxa, and thus contribute to conservation of this ecologically significant group.

## Acknowledgements

We thank Jane Grimwood’s lab at Hudson Alpha for library preparation and sequencing for the Omni-C and PacBio HiFi libraries. This work was done with funding from the University of Georgia, United States National Science Foundation grants DEB-2110875 and DEB-2225028 to J.L.-M., and with funding from the Society for the Study of Evolution’s Rosemary Grant Advanced Award (awarded to E.B.).

## Author Contributions

E.B. conceived of the study with help in framing from J.L.-M.. E.B. performed all bioinformatic analyses, and wrote the original draft. W.R. procured *S. rosea* and *S. psittacina* plants, performed the cross to generate the F1, and maintained plants in the greenhouse. W.R. and J.L.-M. reviewed and edited the manuscript. J.L.-M. and E.B. procured funding for the project.

## Data Availability Statement

Genome assemblies and all sequencing data are deposited into NCBI GenBank and are available under BioProject PRJNA1157763. Genome assemblies and annotations are also available on Zenodo (https://doi.org/10.5281/zenodo.15481882). All code used for assembly and analysis is available at https://github.com/ethan-baldwin/sarracenia_genomics.

## Notes

https://doi.org/10.5281/zenodo.15481882

https://github.com/ethan-baldwin/sarracenia_genomics

### Competing Interest Statement

The authors have declared no competing interest.

## LITERATURE CITED

Albert, V. A., S. E. Williams and M. W. Chase. 1992. Carnivorous plants: phylogeny and structural evolution. Science 257: 1491–1495.

Baldwin, E., M. McNair and J. Leebens-Mack. 2023. Rampant chloroplast capture in Sarracenia revealed by plastome phylogeny. Frontiers in Plant Science 14.

Cai, L., B. J. Arnold, Z. Xi, D. E. Khost, N. Patel, C. B. Hartmann, S. Manickam, et al. 2021. Deeply Altered Genome Architecture in the Endoparasitic Flowering Plant <em>Sapria himalayana</em> Griff. (Rafflesiaceae). Current Biology 31: 1002-1011.e1009.

Cheng, H., G. T. Concepcion, X. Feng, H. Zhang and H. Li. 2021. Haplotype-resolved de novo assembly using phased assembly graphs with hifiasm. Nature Methods 18: 170–175.

Dudchenko, O., M. S. Shamim, S. S. Batra, N. C. Durand, N. T. Musial, R. Mostofa, M. Pham, et al. 2018. The Juicebox Assembly Tools module facilitates <em>de novo</em> assembly of mammalian genomes with chromosome-length scaffolds for under $1000. bioRxiv: 254797.

Durand, N. C., J. T. Robinson, M. S. Shamim, I. Machol, J. P. Mesirov, E. S. Lander and E. L. Aiden. 2016. Juicebox Provides a Visualization System for Hi-C Contact Maps with Unlimited Zoom. Cell Systems 3: 99–101.

Ellison, A. M., N. J. Gotelli, J. S. Brewer, D. L. Cochran-Stafira, J. M. Kneitel, T. E. Miller, A. C. Worley and R. Zamora. 2003. The evolutionary ecology of carnivorous plants. In Advances in Ecological Research. Academic Press. 33: 1–74.

Emms, D. M. and S. Kelly. 2019. OrthoFinder: phylogenetic orthology inference for comparative genomics. Genome Biology 20: 238.

Fasbender, L., D. Maurer, J. Kreuzwieser, I. Kreuzer, W. X. Schulze, J. Kruse, D. Becker, et al. 2017. The carnivorous Venus flytrap uses prey-derived amino acid carbon to fuel respiration. New Phytologist 214: 597–606.

Faust, G. G. and I. M. Hall. 2014. SAMBLASTER: fast duplicate marking and structural variant read extraction. Bioinformatics (Oxford, England) 30: 2503–2505.

Fleischmann, A., J. Schlauer, S. A. Smith and T. J. Givnish. 2017. Evolution of carnivory in angiosperms. In Carnivorous Plants: Physiology, ecology, and evolution. A. Ellison and L. Adamec. Oxford University Press: 0.

Fu, C.-N., S. Wicke, A.-D. Zhu, D.-Z. Li and L.-M. Gao. 2023. Distinctive plastome evolution in carnivorous angiosperms. BMC Plant Biology 23: 660.

Fukushima, K., X. Fang, D. Alvarez-Ponce, H. Cai, L. Carretero-Paulet, C. Chen, T.-H. Chang, et al. 2017. Genome of the pitcher plant Cephalotus reveals genetic changes associated with carnivory. Nature Ecology & Evolution 1: 0059.

Furches, M. S., R. L. Small and A. Furches. 2013. Hybridization leads to interspecific gene flow in Sarracenia (Sarraceniaceae). Am J Bot 100: 2085–2091.

Gabriel, L., T. Brůna, K. J. Hoff, M. Ebel, A. Lomsadze, M. Borodovsky and M. Stanke. 2024. BRAKER3: Fully automated genome annotation using RNA-seq and protein evidence with GeneMark-ETP, AUGUSTUS and TSEBRA. bioRxiv.

Glenn, T. C., R. A. Nilsen, T. J. Kieran, J. G. Sanders, N. J. Bayona-Vásquez, J. W. Finger, T. W. Pierson, et al. 2019. Adapterama I: universal stubs and primers for 384 unique dual-indexed or 147,456 combinatorially-indexed Illumina libraries (iTru & iNext). PeerJ 7: e7755.

Graham, S. W., V. K. Y. Lam and V. S. F. T. Merckx. 2017. Plastomes on the edge: the evolutionary breakdown of mycoheterotroph plastid genomes. New Phytologist 214: 48–55.

Hartmann, S., M. Preick, S. Abelt, A. Scheffel and M. Hofreiter. 2020. Annotated genome sequences of the carnivorous plant Roridula gorgonias and a non-carnivorous relative, Clethra arborea. BMC Res Notes 13: 426.

Harvey, E. and T. E. Miller. 1996. Variance in composition of inquiline communities in leaves of Sarracenia purpurea L. on multiple spatial scales. Oecologia 108: 562–566.

Ibarra-Laclette, E., E. Lyons, G. Hernández-Guzmán, C. A. Pérez-Torres, L. Carretero-Paulet, T.-H. Chang, T. Lan, et al. 2013. Architecture and evolution of a minute plant genome. Nature 498: 94–98.

Kim, D., J. M. Paggi, C. Park, C. Bennett and S. L. Salzberg. 2019. Graph-based genome alignment and genotyping with HISAT2 and HISAT-genotype. Nature Biotechnology 37: 907–915.

Koopman, M. M. and B. C. Carstens. 2010. Conservation genetic inferences in the carnivorous pitcher plant Sarracenia alata (Sarraceniaceae). Conservation Genetics 11: 2027–2038.

Koren, S., A. Rhie, B. P. Walenz, A. T. Dilthey, D. M. Bickhart, S. B. Kingan, S. Hiendleder, et al. 2018. De novo assembly of haplotype-resolved genomes with trio binning. Nature Biotechnology 36: 1174–1182.

Kruse, J., P. Gao, M. Eibelmeier, S. Alfarraj and H. Rennenberg. 2017. Dynamics of amino acid redistribution in the carnivorous Venus flytrap (Dionaea muscipula) after digestion of 13C/15N-labelled prey. Plant Biology 19: 886–895.

Kruse, J., P. Gao, A. Honsel, J. Kreuzwieser, T. Burzlaff, S. Alfarraj, R. Hedrich and H. Rennenberg. 2014. Strategy of nitrogen acquisition and utilization by carnivorous Dionaea muscipula. Oecologia 174: 839–851.

Kuznetsov, D., F. Tegenfeldt, M. Manni, M. Seppey, M. Berkeley, Evgenia V. Kriventseva and E. M. Zdobnov. 2022. OrthoDB v11: annotation of orthologs in the widest sampling of organismal diversity. Nucleic Acids Research 51: D445–D451.

Lan, T., T. Renner, E. Ibarra-Laclette, K. M. Farr, T.-H. Chang, S. A. Cervantes-Pérez, C. Zheng, et al. 2017. Long-read sequencing uncovers the adaptive topography of a carnivorous plant genome. Proceedings of the National Academy of Sciences 114: E4435–E4441.

Li, H. and R. Durbin. 2009. Fast and accurate short read alignment with Burrows-Wheeler transform. Bioinformatics (Oxford, England) 25: 1754–1760.

Li, H., B. Handsaker, A. Wysoker, T. Fennell, J. Ruan, N. Homer, G. Marth, et al. 2009. The Sequence Alignment/Map format and SAMtools. Bioinformatics (Oxford, England) 25: 2078–2079.

Lin, Q., S. S. Yin, M. Mata-Rosas, E. Ibarra-Laclette and T. Renner. 2025. Are carnivorous plants mixotrophic? New Phytol 247: 445–449.

Luciano, C. S. and S. J. Newell. 2017. Effects of prey, pitcher age, and microbes on acid phosphatase activity in fluid from pitchers of Sarracenia purpurea (Sarraceniaceae). PloS one 12: e0181252.

Magallón, S., S. Gómez-Acevedo, L. L. Sánchez-Reyes and T. Hernández-Hernández. 2015. A metacalibrated time-tree documents the early rise of flowering plant phylogenetic diversity. New Phytol 207: 437–453.

Malmberg, R. L., W. L. Rogers and M. S. Alabady. 2018. A carnivorous plant genetic map: pitcher/insect-capture QTL on a genetic linkage map of Sarracenia. Life Sci Alliance 1: e201800146.

Mendes, F. K., D. Vanderpool, B. Fulton and M. W. Hahn. 2020. CAFE 5 models variation in evolutionary rates among gene families. Bioinformatics (Oxford, England) 36: 5516–5518.

Neumann, P., A. Navrátilová, A. Koblížková, E. Kejnovský, E. Hřibová, R. Hobza, A. Widmer, et al. 2011. Plant centromeric retrotransposons: a structural and cytogenetic perspective. Mob DNA 2: 4.

Nevill, P. G., K. A. Howell, A. T. Cross, A. V. Williams, X. Zhong, J. Tonti-Filippini, L. M. Boykin, et al. 2019. Plastome-Wide Rearrangements and Gene Losses in Carnivorous Droseraceae. Genome Biology and Evolution 11: 472–485.

Nguyen, L. T., H. A. Schmidt, A. von Haeseler and B. Q. Minh. 2015. IQ-TREE: a fast and effective stochastic algorithm for estimating maximum-likelihood phylogenies. Mol Biol Evol 32: 268–274.

Paajanen, P., G. Kettleborough, E. López-Girona, M. Giolai, D. Heavens, D. Baker, A. Lister, et al. 2019. A critical comparison of technologies for a plant genome sequencing project. GigaScience 8.

Palfalvi, G., T. Hackl, N. Terhoeven, T. F. Shibata, T. Nishiyama, M. Ankenbrand, D. Becker, et al. 2020. Genomes of the Venus Flytrap and Close Relatives Unveil the Roots of Plant Carnivory. Current Biology 30: 2312-2320.e2315.

Pavlovič, A. 2022. Photosynthesis in Carnivorous Plants: From Genes to Gas Exchange of Green Hunters. Critical Reviews in Plant Sciences 41: 305–320.

Rentsch, J. D. and R. C. Holland. 2020. Population Genetic Structure and Natural Establishment of Hybrids Between Sarracenia flava and Sarracenia minor in Francis Marion National Forest. Castanea 85: 108-121, 114.

Sigman, M. J. and R. K. Slotkin. 2016. The First Rule of Plant Transposable Element Silencing: Location, Location, Location. Plant Cell 28: 304–313.

Simão, F. A., R. M. Waterhouse, P. Ioannidis, E. V. Kriventseva and E. M. Zdobnov. 2015. BUSCO: assessing genome assembly and annotation completeness with single-copy orthologs. Bioinformatics (Oxford, England) 31: 3210–3212.

Stephens, J. D. and D. R. Folkerts. 2012. Life History Aspects of <i>Exyra semicrocea</i> (Pitcher Plant Moth) (Lepidoptera: Noctuidae). Southeastern Naturalist 11: 111-126, 116.

Sun, G., Y. Xu, H. Liu, T. Sun, J. Zhang, C. Hettenhausen, G. Shen, et al. 2018. Large-scale gene losses underlie the genome evolution of parasitic plant Cuscuta australis. Nature Communications 9: 2683.

Veleba, A., F. Zedek, L. Horová, P. Veselý, M. Srba, P. Šmarda and P. Bureš. 2020. Is the evolution of carnivory connected with genome size reduction? American Journal of Botany 107: 1253–1259.

Wicke, S., B. Schäferhoff, C. W. dePamphilis and K. F. Müller. 2013. Disproportional Plastome-Wide Increase of Substitution Rates and Relaxed Purifying Selection in Genes of Carnivorous Lentibulariaceae. Molecular Biology and Evolution 31: 529–545.

Yu, G., L. G. Wang, Y. Han and Q. Y. He. 2012. clusterProfiler: an R package for comparing biological themes among gene clusters. Omics 16: 284–287.

Zhou, C., S. A. McCarthy and R. Durbin. 2023. YaHS: yet another Hi-C scaffolding tool. Bioinformatics (Oxford, England) 39: btac808.

Zhu, Q.-g., Y. Xu, Y. Yang, C.-f. Guan, Q.-y. Zhang, J.-w. Huang, D. Grierson, et al. 2019. The persimmon (Diospyros oleifera Cheng) genome provides new insights into the inheritance of astringency and ancestral evolution. Horticulture Research 6: 138.

